# Scalable fold-change estimation from microbial metagenomics

**DOI:** 10.64898/2026.10.08.757531

**Authors:** Sarah Teichman, Amy D Willis

**Affiliations:** Department of Biostatistics, University of Washington

**Keywords:** microbiome, regression, compositional data, differential abundance, genomics, algorithms

## Abstract

Microbial sequencing data are subject to sample-specific and species-specific distortions that complicate inference about true microbial abundances. Recent work established assumptions under which relative fold-differences in microbial abundances can be estimated from sequencing data, accompanied by an estimation and testing method called `radEmu`. While `radEmu` has excellent error rates, its computational cost can be prohibitive for large-scale analyses. Here we introduce `fastEmu`, a scalable approximation to `radEmu`’s testing methodology. `fastEmu` uses a small, data-driven subset of taxa to approximate the computationally expensive test statistic. In a metagenomic analysis, we show a reduction in per-taxon runtime from 4-8 hours to 0.5-45 seconds, and simulations illustrate no adverse impact on estimation error or Type I error rate control. `fastEmu` enables testing differential abundance for thousands of microbial taxa while retaining clear parameter interpretation and error rate control.

---

Microbial metagenomics, including shotgun sequencing and amplicon sequencing, can be used to construct count or coverage data on microbial genomes and genes. It is widely appreciated that both sample-specific (e.g., sequencing depth) and species-specific (e.g., DNA extraction efficiency) effects make it implausible that these data directly reflect microbial abundances. To formalize what can and cannot be learned about environmental abundances from metagenomic data, Clausen, Teichman and Willis (2024) studied assumptions under which important summaries of microbial abundances can be learned from sequencing data. A key result is that under the assumption that sequencing counts are equal (on average) to the product of true microbial abundances with sample-specific effects and species-specific effects, fold-differences in average abundances *cannot* be estimated from sequencing data. In contrast, *relative* fold-differences *can* be estimated, where relative refers to either a reference group (a species known to be unchanging in abundance, such as a spiked-in taxon) or the average fold-difference across all categories. In addition to their identifiability results, Clausen, Teichman and Willis (2024) develop a method (“radEmu”) to estimate relative fold-differences in true microbial abundances from sequencing data. The method also permits hypothesis testing that fold-differences are equal to typical fold-differences (“differential abundance”).

There are two steps in a typical data analysis using `radEmu`. Firstly, the parameters of the model are estimated; that is, numerical values representing the best-fitting difference in absolute abundance is obtained for each species-covariate combination. Secondly, a test of the hypothesis that this fold-difference is equal to the typical fold-differences is performed. These two steps are decoupled because the testing methodology (robust score tests) requires estimating the parameters of the model under the null hypothesis. Unfortunately, fitting under the null hypothesis can be computationally burdensome, as even the fastest algorithms for this problem may converge slowly in practice. Despite their outstanding small sample error rate control, the computational burden of robust score tests can be prohibitive for datasets containing a large number of species, even on a per-species basis. For example, in the analysis of ≈ 8, 000 taxa described below, each score test takes between four and eight hours to complete on a high-performance computing cluster, and 8, 000 tests must be run.

To address the computational limitations of `radEmu`, we propose a new, scalable method for testing hypotheses about fold-differences in microbial abundances. Our approach, called `fastEmu`, tests a hypothesis that is numerically very similar to the `radEmu` null hypothesis, but is substantially faster to test. The aforementioned score tests take at most 45 seconds each with `fastEmu`, with a median runtime of 3 seconds.

We briefly describe the methodology of `fastEmu` and contrast it with `radEmu`. Let *W*_*ij*_ denote the (unknown) true microbial abundance or concentration (e.g., in DNA copies per microliter) of species *j*= 1, …, *J* in sample *i*= 1, …, *n*, and let *Y*_*ij*_ denote the (observed) coverage or observed number of copies of species *j* in sample *i* obtained from high-throughput sequencing. We would like to estimate the log fold-differences *β*_*kj*_ in the model 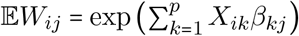 but we observe the *Y*_*ij*_’s, not the *W*_*ij*_’s. The two are linked via E*Y*_*ij*_ = *s*_*i*_ × *e*_*j*_ × E*W*_*ij*_ where E denotes the expected value (average) of a random variable, *X*_*ik*_ denotes the value of the *k*th covariate in sample *i, s*_*i*_ is the unknown sampling effect for sample *i* and *e*_*j*_ is the unknown detection efficiency of species *j* (McLaren, Willis and Callahan, 2019). Clausen, Teichman and Willis (2024) demonstrated that while it is not possible to estimate or test hypotheses about *β*_*kj*_ from 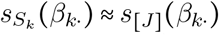, it is possible to estimate and test *β*_*kj*_ − *g*(*β*_*k*⋅_), where *g* is a centering function. They argue that choosing *g* to be the smoothed median (over taxa in [*J* ] := {1, …, *J* }) is an appropriate and interpretable choice for microbiome data analysis, because even if some fold-differences are large, *g (β*_*k*_ ) ≈ 0 if most fold-differences are small. Similarly, if the distribution of fold differences is approximately symmetric, *g (β*_*k*_) ≈ 0. In these circumstances, *β*_*kj*_ − *g (β*_*k*_) ≈*β*_*kj*_ and `radEmu` estimates fold-differences in abundance on the absolute scale.

The core innovation of `fastEmu` is that the computationally expensive *β*_*kj*_ −*g*(*β*_*k*⋅_) can be replaced by a computationally cheap approximation. For a set *S* ⊆ [*J* ] and a *J* –dimensional vector *b*, let *s*_*S*_(*b*) denote the smoothed median over the elements of *b* with indices in *S*. `fastEmu` finds small subsets *S*_*k*_ of [*J* ] such that 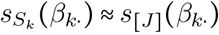, and uses them to test the approximate null hypothesis 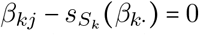. Our approach involves using the data to select a hypothesis that is then tested using the same data. Our simulation study compares the impact of this “double dipping” approach to a sample-splitting approach, and we find no negative impacts on estimation error nor error rate control. In contrast to typical doubledipping approaches, `fastEmu` *approximates* a well-defined, pre-specified null hypothesis; it is not attempting to identify hypotheses that are most likely to be rejected. `fastEmu` is also efficient when *g* ( *β*_*k*_⋅)=( *β*_*kj*_ ′)for a fixed reference taxon *j* ′, which is applicable when DNA is spiked-in to samples.

We illustrate the computational feasibility and statistical advantages of `fastEmu` by investigating how marine surface microbes change in abundance across temperature gradients in temperate oceans. We consider 8, 041 genome bins in 57 samples from the surface and shallow ocean with temperatures between 10 and 30 degrees Celsius using on data collated and organized by Schechter et al. (2025) (see Supplementary Material Section 3). We esti-mated model parameters using `radEmu`, with *Y*_*ij*_’s equal to the 25% trimmed mean coverage (“Q2Q3”) of the ribosomal protein L14, and temperature as the single predictor *X*_*i*_. Fitting the `radEmu` model took 188 minutes on a high-performance computing cluster. Testing the null hypothesis that there is no difference in the abundance of genome bin *j* across oceans differing in temperature (relative to the typical difference across genome bins) took between 0.5and 45 seconds per genome bin using `fastEmu`. For comparison, we tested the same hypotheses for a random 5% of genome bins, and these took between 4.5 and 8 hours using `radEmu`.

The approximation that accompanies this >7, 000 ×speed-up is miniscule: exact (`radEmu`) and approximate (`fastEmu`) parameter estimates differ by 9 × 10^−5^, which is orders of magnitude below the median absolute effect size of 0.3. While comparator methods remain faster than `fastEmu` (analogous tests took a total of 14 seconds, 8 minutes, 18 minutes, and 34 minutes for LinDA (Zhou et al., 2022), MaAsLin3 (Nickols et al., 2024), ANCOM-BC2 (Lin and Peddada, 2024), and ALDEx2 (Fernandes et al., 2013) respectively), these methods do not control Type 1 error rate in small samples (Clausen, Teichman and Willis, 2024, Figure 2), leading to unreliable p-values in general. Furthermore, these methods generally estimate differences in centered log-ratio transformed sequencing data, with an unclear connection to microbial abundances. Supplementary material for this article includes the `fastEmu` algorithm, runtime and error rate simulations, and additional details (including a detailed cross-method comparison) for the marine surface microbes data analysis.

**Fig 1:**
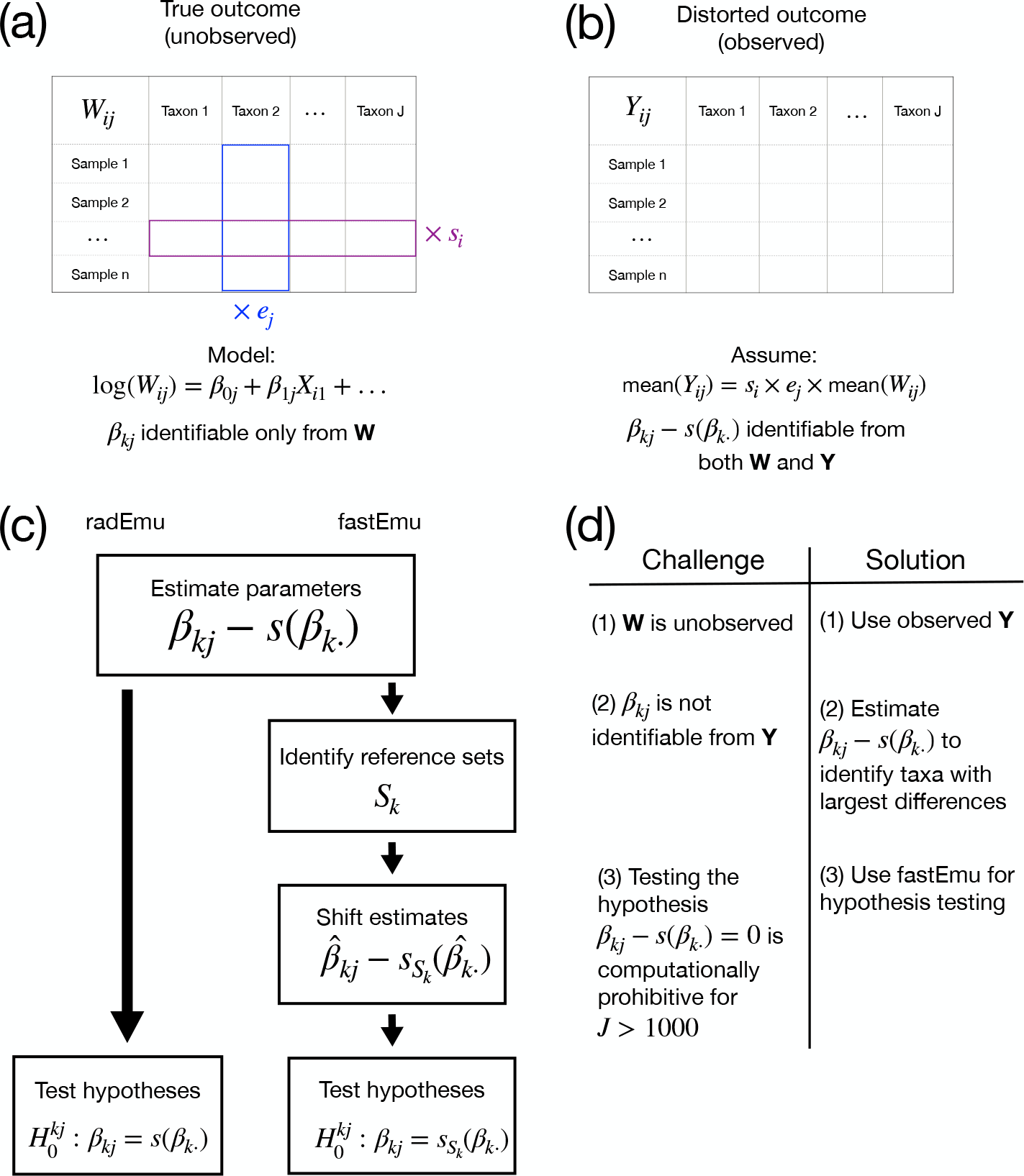
(a) In microbiome studies, true microbial abundances are desired but unobserved. If this information was known, fold-differences in abundances could be estimated using a log-linear model. (b) Instead, noisy and distorted data is observed. Previous literature demonstrated that a centered modification of the desired fold-difference parameters can be learned from the distorted data. (c) The algorithmic steps followed by `radEmu` (Clausen, Teichman and Willis, 2024) and `fastEmu` (this work), and (d) a summary of the statistical and computational challenges in microbial abundance fold-difference estimation.

**Fig 2:**
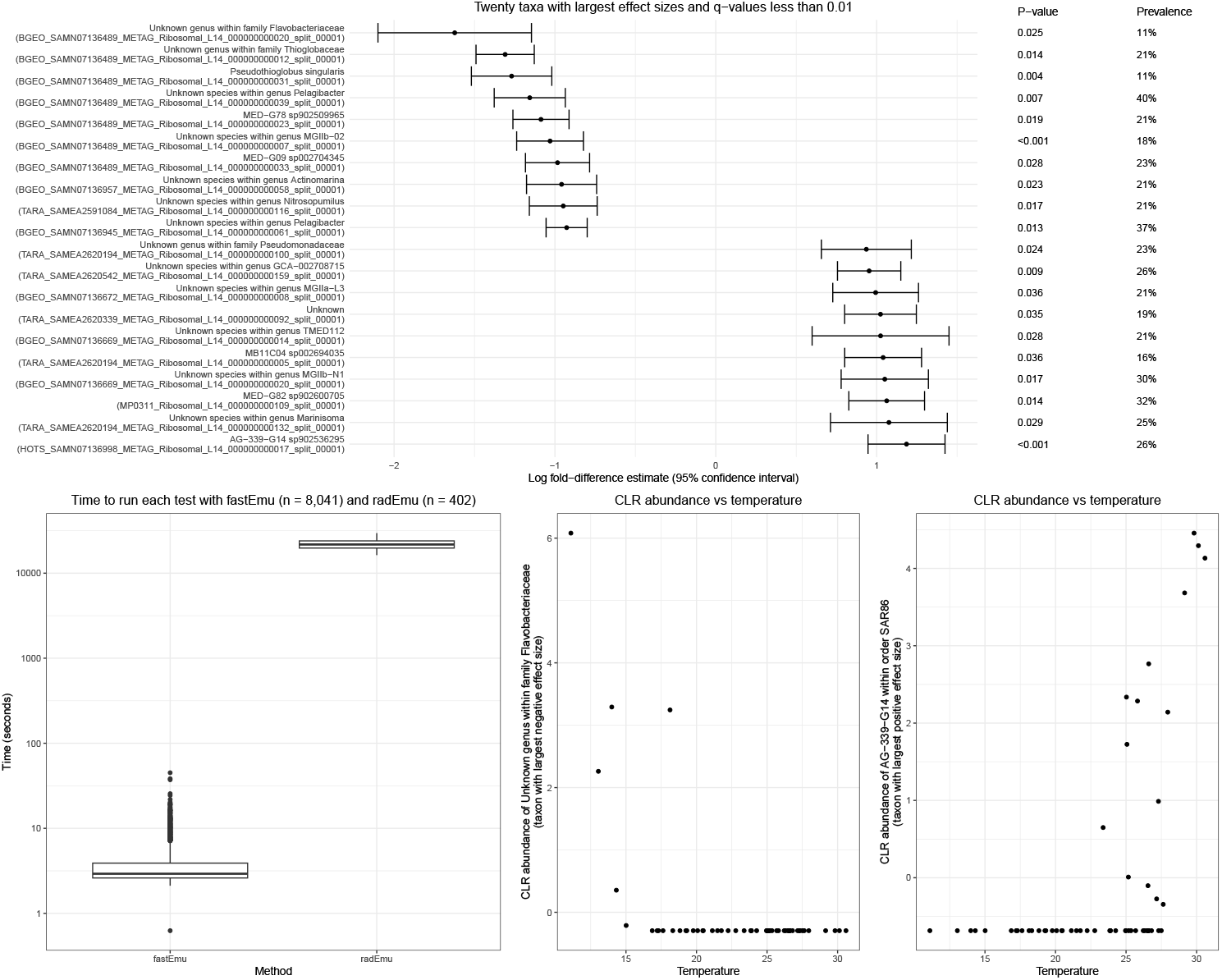
(Upper panel) The 20 taxa from the marine surface microbes dataset with the largest absolute log fold-difference estimates and *q*-values less than 0.01. Taxa are ordered by effect size. Points and horizontal bars show estimated log fold-differences and 95% confidence intervals, respectively. Robust score test p-values and taxon prevalences are reported alongside each taxon. (Lower left panel) Runtime comparison of robust score tests implemented in `fastEmu` and `radEmu`. (Lower middle and right panels) Relationships between CLR-transformed abundances and ocean temperature for the taxa with the largest negative (middle) and positive (right) estimated effects, respectively. Data collated by Schechter et al. (2025).

## Data availability

The datasets analysed during the current study are available at https://api.figshare.com/v2/file/download/53897114. Further details are provided in Supplementary Material Section 3.

## Acknowledgements

We gratefully acknowledge A. Murat Eren and Matthew S. Schechter for their assistance with the marine surface microbes dataset.

## Notes

https://github.com/statdivlab/fastEmu

### Competing Interest Statement

The authors have declared no competing interest.

### Funding

- National Institute of General Medical Sciences: R35 GM133420

