## Supplementary Material for "Scalable fold-change estimation from microbial metagenomics"

### 1. Algorithms.

1.1. *Smoothed median.* As in the main text, we use $s_S(b)$ to denote the smoothed median over the elements of $b$ with indices in $S$. The smoothed median is defined as follows:

$$(1) \quad s_S(b) = \operatorname{argmin}_c \sum_{j \in S} h_\delta(b_j - c)$$

$$(2) \quad h_\delta(x) = \delta^2 \left( \sqrt{1 + (x/\delta)^2} - 1 \right)$$

for $b_j$ defined to be the $j$th element of $b$, and a fixed parameter $\delta > 0$ that controls the “smoothness” of the smoothed median. As $\delta \downarrow 0$, $s(b)$ approximates the median of $b$, and as $\delta \rightarrow \infty$, $s(b)$ approximates the mean of $b$. We choose $\delta = 0.1$ throughout.

1.2. *fastEmu algorithm.* The fastEmu algorithm makes use of two radEmu algorithms: (a) the iterative maximum likelihood algorithm, which estimates the parameters under the alternative hypothesis (Clausen, Teichman and Willis (2024, Section 3.1)), and (b) the augmented Lagrangian algorithm, which estimates the parameters under the null hypothesis (Clausen, Teichman and Willis (2024, Section 3.2)). We refer to the former algorithm as the “$H_A$ algorithm” and the latter as the “$H_0$ algorithm”, reflecting their emphases on estimation under the alternative ($H_A$) and null ($H_0$) hypothesis.

The fastEmu algorithm is as follows:

1. Estimate parameters $\beta_{kj} - s_{[J]}(\beta_{k\cdot})$ for each taxon $j \in [J]$ and each covariate $k \in \{0, \dots, p-1\}$ using the $H_A$ algorithm.
2. For a prespecified reference set size $m$, for each $k \in \{0, \dots, p-1\}$, identify the $m$ taxa with the smallest values of $|\hat{\beta}_{kj} - s_{[J]}(\hat{\beta}_{k\cdot})|$. Call this data-driven reference set $S_k \subset [J]$.
3. For each taxon $j \in \{1, \dots, J\}$ and each covariate $k \in \{1, \dots, p-1\}$ associated with a parameter to be tested:
  a) Define $S_j = (\cup_{k \in \{0, \dots, p-1\}} S_k) \cup \{j\}$.
  b) Define $Y_j^\dagger$ to be the $n \times |S_j|$ submatrix of $Y$ containing column indices $j' \in S_j$. Define $\beta_{kj}^\dagger$ to be the true log fold-differences of the mean of category $j \in S_j$ across subpopulations with unit differences in covariate $k$ but otherwise matched on covariates. Use the $H_0$ algorithm applied to data $Y^\dagger$ to estimate $\beta_{k'j'}^\dagger - s_{S_{k'}}(\beta_{k'j'}^\dagger)$ for each $j' \in S_j$ and $k' \in \{0, \dots, p-1\}$ under the null hypothesis $\beta_{kj}^\dagger - s_{S_k}(\beta_{k\cdot}^\dagger) = 0$. Call the estimates $\hat{\beta}_{k'j'}^\dagger - s_{S_{k'}}(\hat{\beta}_{k'j'}^\dagger)$.
  c) Calculate the Hessian, score, and constraint derivative at $\hat{\beta}_{k'j'}^\dagger - s_{S_{k'}}(\hat{\beta}_{k'j'}^\dagger)$ to compute the robust score test statistic using the construction of $T_{RS}$ in Section 5 of Clausen, Teichman and Willis (2024).

We prove in Section 5 that given a **pre-specified reference set**, fastEmu and radEmu target the same parameters and that the fastEmu test statistic has the correct asymptotic distribution. We also prove that the approach of using `radEmu` estimates to determine **data-driven reference sets** $S_k$ has negligible asymptotic impacts. We compare estimates, test results, and test timing through simulation and a data analysis in Section 2 and Section 3.

**2. Simulations.** We conduct a simulation study to compare the computation time, empirical type 1 error rate, and empirical power of our proposed test compared to the robust score test from `radEmu` Clausen, Teichman and Willis (2024). In Clausen, Teichman and Willis (2024) the authors compared `radEmu` to several popular differential abundance methods, including ALDEx2 (Fernandes et al., 2013), ANCOM-BC2 (Lin and Peddada, 2024), LinDA (Zhou et al., 2022), and MaAsLin3 (Nickols et al., 2024). We refer the reader to those simulation studies for a comprehensive comparison of the empirical type 1 error rates and power of these methods across a range of simulation settings. In this paper, we constrain our simulation study to our proposed test and the robust score test implemented in `radEmu` (Clausen, Teichman and Willis, 2024).

**2.1. Simulation setup.** We use a similar simulation setup to Clausen, Teichman and Willis (2024). First, define the true unknown reference set $S_k^*$ as the set of $m$ taxa with the smallest values of $\beta_{kj} - s_{[J]}(\beta_{k\cdot})$. We test the hypothesis $\beta_{k^*j^*} - s_{[J]}(\beta_{k\cdot}) = 0$ for `radEmu`, and the hypothesis $\beta_{k^*j^*} - s_{S_k^*}(\beta_{k\cdot}) = 0$ for `fastEmu`, for $k^* = 2$ and $j^* = J/2$. We design our simulation such that $s_{[J]}(\beta_{k\cdot}) = s_{S_k^*}(\beta_{k\cdot})$, in order that the same hypothesis is tested by both methods, thereby producing a fair comparison.

We fix $p = 3$ and vary $n \in \{20, 50, 250\}$ and $J \in \{250, 500, 1000, 2000, 3000, 4000, 5000\}$. We generate data from a zero-inflated negative binomial (ZINB) distribution. For all $i$, we set $X_{i0} = 1$ and draw $X_{i1} \sim \text{Uniform}(0, 1)$ and $X_{i2} \sim \text{Normal}(0, 2)$ such that $X_{i1}$ and $X_{i2}$ are independent of each other. $\beta_0$, the first row of $\beta$, has odd elements forming a linear sequence from $-3$ to $3$ and even elements forming a linear sequence from $3$ to $-3$. $\beta_1$ is given by $5 \times \sinh(x)/\sinh(10)$ where $x$ is a linearly sequence from $-10$ to $10$ ($\sinh$ is the hyperbolic sine function). We set the $J/2$ element to $b$, where $b = 0$ in simulations under the null (to assess Type 1 error) and $b \in \{1, 2, 3, 4, 5\}$ in simulations under the alternative (to assess power). $\beta_2$ is a random permutation of $\beta_1$. We generate taxon effects $\delta$ as a random permutation of $J$ evenly spaced values within the range $[-1, 1]$. Sample effects are drawn with $z_i \sim N(\log(50/r_i), 1)$ with $r_i = 1/J \sum_j \exp(X_i \beta^j + \delta_j)$, giving a constant expected sequencing depth $\sum_j \mathbb{E}[Y_{ij}]$ across samples, despite variation in $z_i$.

We construct $\beta$, $\delta$ and $\mathbf{z}$ in this way such that $|\beta_{k^*j^*} - s_{[J]}(\beta_{k\cdot})| \approx b$ for $j^* = J/2$, with exact equality when $b = 0$. Most categories have a similar log fold-difference in abundance across levels of $X_{i1}$ and $X_{i2}$, and a few categories have log fold-differences with large magnitudes. The values of $b$ under the alternative are intended to reflect effect sizes ranging from a $\approx 1.7$-fold to a 150-fold deviation from the typical category’s log-fold difference across unit differences in $X_{i1}$. We run 500 trials for each simulation setting. Score tests were run within serial simulation tasks run on a single CPU core of a high-performance computing cluster, with memory allocations ranging from 2 to 36 GB depending on the simulation setting.

**2.2. Time comparison.** We compare the timing of `fastEmu` robust score tests to the timing of `radEmu` robust score tests in Figure 1. This figure shows that the median time to run `radEmu` score tests increases superlinearly with $J$. For a fixed $J$, the interquartile range of time for these tests are stable across sample sizes, although the maximum runtimes are higher for smaller sample sizes. The time to run `fastEmu` score tests increases slightly with $n$ and $J$, but the median test time ranges between 20 and 70 seconds across $n$ and $J$ settings. In these simulation settings, aggregating over sample size, `fastEmu` is only 2 and 5 times faster on average than `radEmu` for smaller $J$ of 250 and 500 respectively. However, for larger $J$ of 3000 and 4000, `fastEmu` is 152 and 265 times faster than `radEmu` on average.

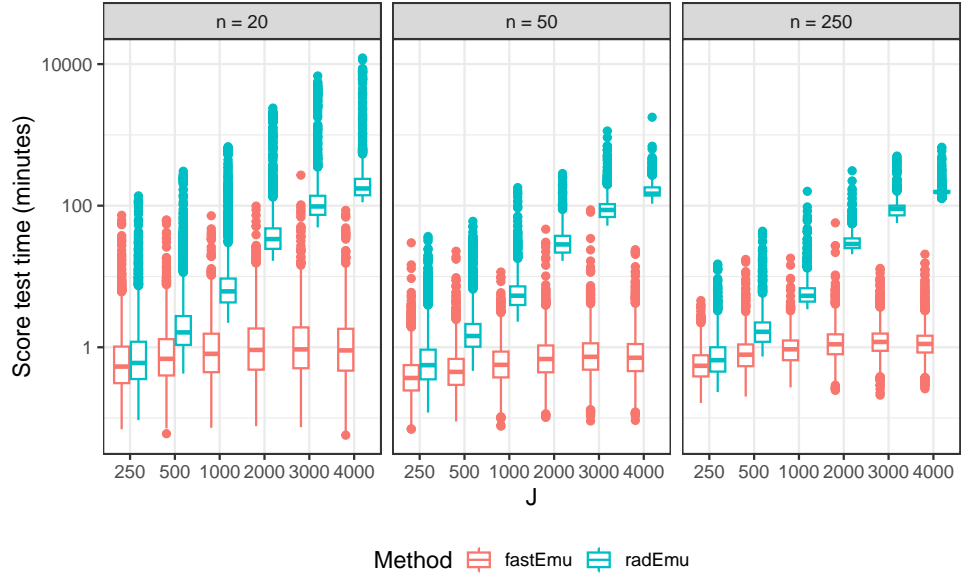

Fig 1: Boxplots of the time to run a single score test within simulation settings across different values of $n$ and $J$. Data are simulated using the model in Equation (2) and draws from a zero-inflated Negative Binomial (ZINB) distribution. P-values are generated from robust score tests from full and reduced models.

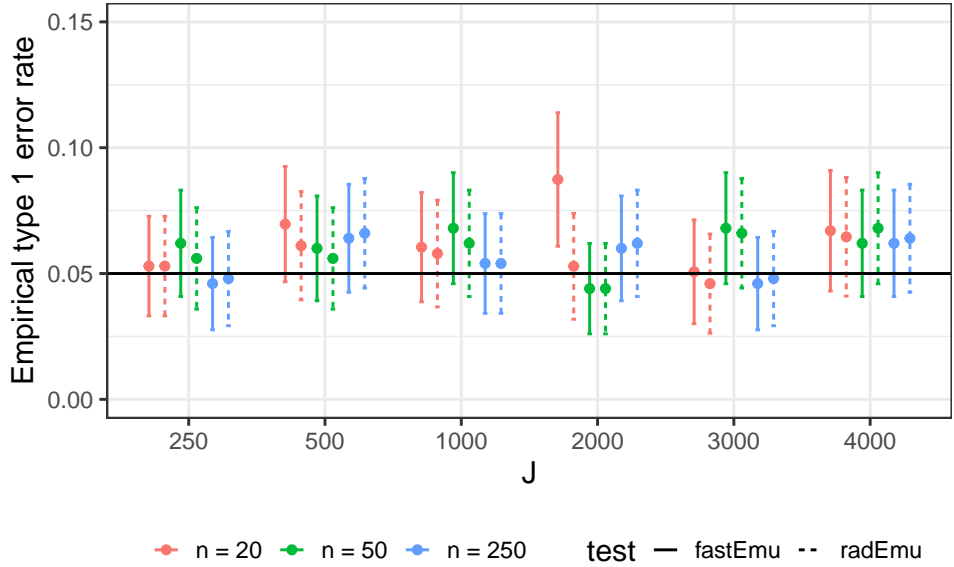

Fig 2: Empirical type 1 error rates for robust score tests using an alpha level of 5% across 500 trials. 95% confidence intervals for the true error rates are shown. Data are simulated using the model in Equation (2) and draws from a zero-inflated Negative Binomial distributions. P-values are generated from robust score tests using the full models (radEmu) and reduced models (fastEmu).

**2.3. Type 1 error rate control comparison.** We now compare the Type 1 error rates of `radEmu` and `fastEmu`. Figure 2 presents empirical Type 1 error rates for a test performed at a nominal level of 5%. We also show 95% confidence intervals for the true error rates based on 500 trials for each simulation setting. Each confidence interval contains the nominal rate of 5%, except for `fastEmu` tests in the simulation setting with $J = 2000$ and $n = 20$. We believe that this is due to ill-conditioning of the observed information matrix (discussed further in Section 2.5) due to Monte Carlo variation, as ill-conditioning is much more likely at small sample sizes. Overall, we see that these two methods have comparable type 1 error rate control across a range of sample sizes and $J$. This also demonstrates that both methods have an empirical type 1 error rate control that is close to the nominal level of 0.05 for each of the sample sizes tested.

**2.4. Power comparison.** Finally, we compare the power of `fastEmu` to `radEmu` in our simulated data. Figure 3 shows that `fastEmu` attains the same or higher power as `radEmu`. `fastEmu` tends to have higher power than `radEmu` for large signal magnitudes, smaller sample sizes, and larger $J$. It is not surprising that these methods have slightly different power as they test slightly different hypotheses. These results imply that for a given sample size and signal magnitude, the ability of `radEmu` to detect that signal decreases with increasing $J$, while `fastEmu` has more consistent power with increasing $J$.

**2.5. Estimation failures.** In each setting with a sample size of 20, there are a subset of trials that failed during the parameter estimation step of `radEmu` (preventing computation of the subsequent robust score tests in both `radEmu` and `fastEmu`). This failure is caused by a non-positive definite observed information matrix, preventing evaluation of the Firth penalty used during parameter estimation (Clausen, Teichman and Willis, 2024). The Firth penalty is included to prevent parameter estimates from diverging under sparse data, separation or quasi-separation. The number of failures increases with $J$, with approximately 2%, 4%, 7%, 12%, 14%, and 16% failures for $J \in \{250, 500, 1000, 2000, 3000, 4000\}$ respectively. We exclude situations where estimation cannot be performed from our empirical error rates calculations.

**3. Differential abundance of marine microbes across temperature gradients.** Here we study the abundance of marine surface microbes across gradients of temperature using genomes from Paoli et al. (2022); Martínez-Pérez et al. (2022); Delmont and Eren (2018); Delmont et al. (2019) as collated and organized into genome bins by Schechter et al. (2025) using the EcoPhylo workflow of anvio Eren et al. (2015, 2021). After following the instructions in the reproducible workflow of Schechter et al. (2025) available at https://merenlab.org/data/ecophylo-ribosomal-proteins/#ecophylo-interactive-interface-of-the-open-surface-ocean, we used the anvio program `anvi-export-table` to export 25% trimmed mean coverage values (“Q2Q3”) from the profile database for ribosomal protein L14. We subset to the $n = 57$ samples collected from the surfaces of oceans with temperatures between 10 and 30 degrees Celsius, thereby excluding samples from the Arctic and Southern Oceans. We exclude these colder oceans in order to understand how the abundance of surface-dwelling microbes in moderate to warm oceans will respond to changes in temperature. These data give rise to our analysis of the abundance of 8,041 genome bins. For simplicity, we refer to these genome bins as “taxa.”

We perform differential abundance analysis with `fastEmu`, and compare results to those from ANCOM-BC2 (Lin and Peddada, 2024), ALDEx2 (Fernandes et al., 2013), LinDA (Zhou et al., 2022), and MaAsLin3 (Nickols et al., 2024). We also perform the computationally expensive `radEmu` robust score tests for a subset of taxa.

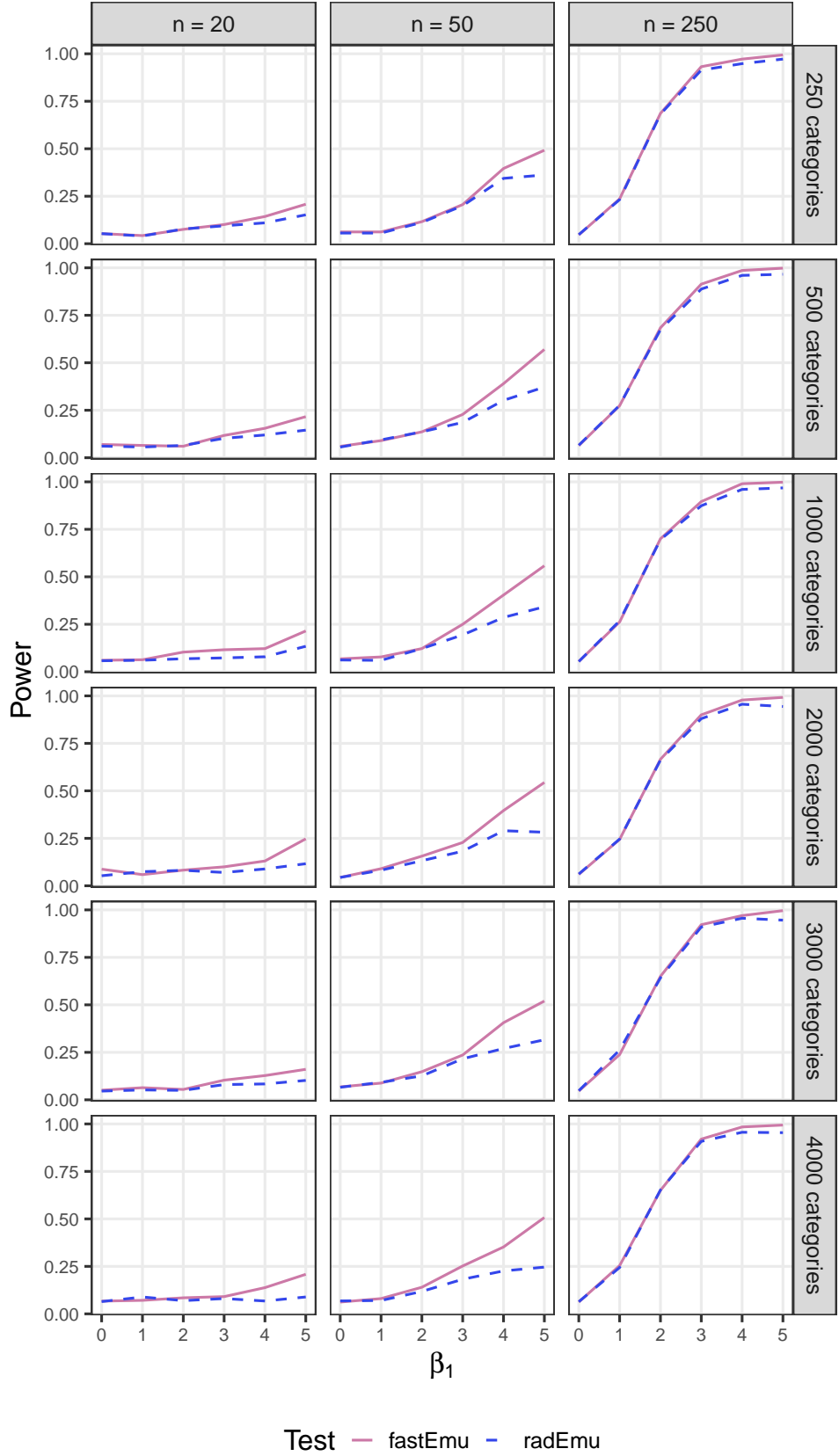

Fig 3: Empirical power for robust score tests using an alpha level of 5%, across 500 trials. Data are simulated using the model in Equation (2) and draws from a zero-inflated Negative Binomial distributions, under specific alternative hypotheses values that range from 1 to 5. P-values are generated from robust score tests using the full models (radEmu) and reduced models (fastEmu).

**3.1. Time comparison.** Parameter estimation for `radEmu` (a necessary precursor to `fastEmu`) took 188 minutes. Each `fastEmu` robust score test took between 0.5 and 45 seconds. For comparison, we ran `radEmu` robust score tests for 5% of taxa, and they each took between 4.5 and 8 hours to complete. On average, for the taxa for which we ran `radEmu` robust score tests, the `fastEmu` tests were 7,034 times faster. The comparator methods all have shorter runtimes than `radEmu` and `fastEmu`, estimating parameters and testing hypotheses for all 8,041 taxa in 14 seconds, 8 minutes, 18 minutes, and 34 minutes for `LinDA`, `MaAsLin3`, `ANCOM-BC2`, and `ALDEx2` respectively. While `fastEmu` still takes longer to run than other differential abundance methods, the score tests are fast and easily parallelizable. Importantly, we note that metagenomic assembly and genome recovery for even a subset of the data considered here (Paoli et al., 2022) required weeks of high-performance computing time; and therefore interpretable differential abundance analysis with Type 1 error rate control contributes very modestly to the computational expensive of analyzing these data.

**3.2. Comparison of estimates.** There are no missing estimates from `radEmu`, `fastEmu`, or `LinDA`. `ANCOM-BC2` and `MaAsLin3` have missing estimates for the 240 taxa that appear only in a single sample in the dataset. `ALDEx2` has 285 missing estimates: 120 for taxa that appear only in a single sample, and the remaining 165 from other taxa with low prevalence.

In Figure 4, we compare log fold-difference estimates across each differential abundance method that we consider. Log fold-difference estimates from `radEmu` (not shown) and `fastEmu` are nearly identical. By construction, they are equal up to a constant shift of $|s_{[J]}(\hat{\beta}_{1\cdot}) - s_{S_1}(\hat{\beta}_{1\cdot})|$, which in this analysis is $9 \times 10^{-5}$. Log fold-difference estimates from `ALDEx2` and `LinDA` have a Spearman correlation of 0.99 and estimates from `ANCOM-BC2` and `MaAsLin3` have a Spearman correlation of 0.97. All other Spearman correlations between pairs of methods range between 0.62 and 0.88. `fastEmu` has higher Spearman correlations with `ALDEx2` and `LinDA` than with `ANCOM-BC2` and `MaAsLin3`.

As seen in Figure 4, the estimates from `fastEmu` have a bimodal distribution, with a mode located slightly above 0 and a mode located slightly below 0. An estimate of 0 from `fastEmu` represents the approximate median log fold-difference over all taxa. If we assume that the median fold-difference across these taxa is zero, then this implies that there is a large number of taxa whose abundance is slightly greater in warmer temperatures, and a smaller group of taxa whose abundance slightly lower in warmer temperatures, on average. The estimates from `ANCOM-BC2` and `MaAsLin3` have a more subtle bimodal pattern. The estimates from `ALDEx2` and `LinDA` appear unimodal at 0. However, unlike `fastEmu`, estimates from these methods cannot be directly connected to statements about microbial abundances, and no comparison of implied population structures is possible.

`ANCOM-BC2` and `MaAsLin3` estimates both have large spreads, with tails that extend to estimates with large magnitudes of 34 and 166 respectively. The taxa with estimates in the tails of these distributions are taxa with low prevalence in the dataset. In the truncated histograms in Figure 4, the tails of these distributions are dominated by low prevalence taxa. `ALDEx2` and `LinDA` have the opposite pattern. `ALDEx2` has the smallest spread of estimates (all estimates have magnitudes less than 0.5) and `LinDA` has a moderate spread of estimates (magnitudes of 2). Taxa with low prevalence have estimates that are concentrated near 0, and the tails are made up of taxa with higher prevalence. Finally, `fastEmu` and `radEmu` estimates have a moderate spread, with magnitudes up to 3, and a longer left tail of negative log fold-differences. This left tail is dominated by taxa with low prevalence.

The differences in the behavior of each of these methods for low prevalence taxa is important, as it can be difficult to determine whether a rare taxon is truly differentially abundance across conditions. `ALDEx2` and `LinDA` are both conservative for these taxa, shrinking estimates towards zero. `ANCOM-BC2` and `MaAsLin3` behave in an opposite manner, with large estimates for parameters associated with taxa that appear in only a few samples. `radEmu` and `fastEmu` provide a middle ground, in which many low prevalence taxa appear in the tails of the log fold-difference distribution, but with estimates of smaller magnitudes compared to taxa with higher prevalence.

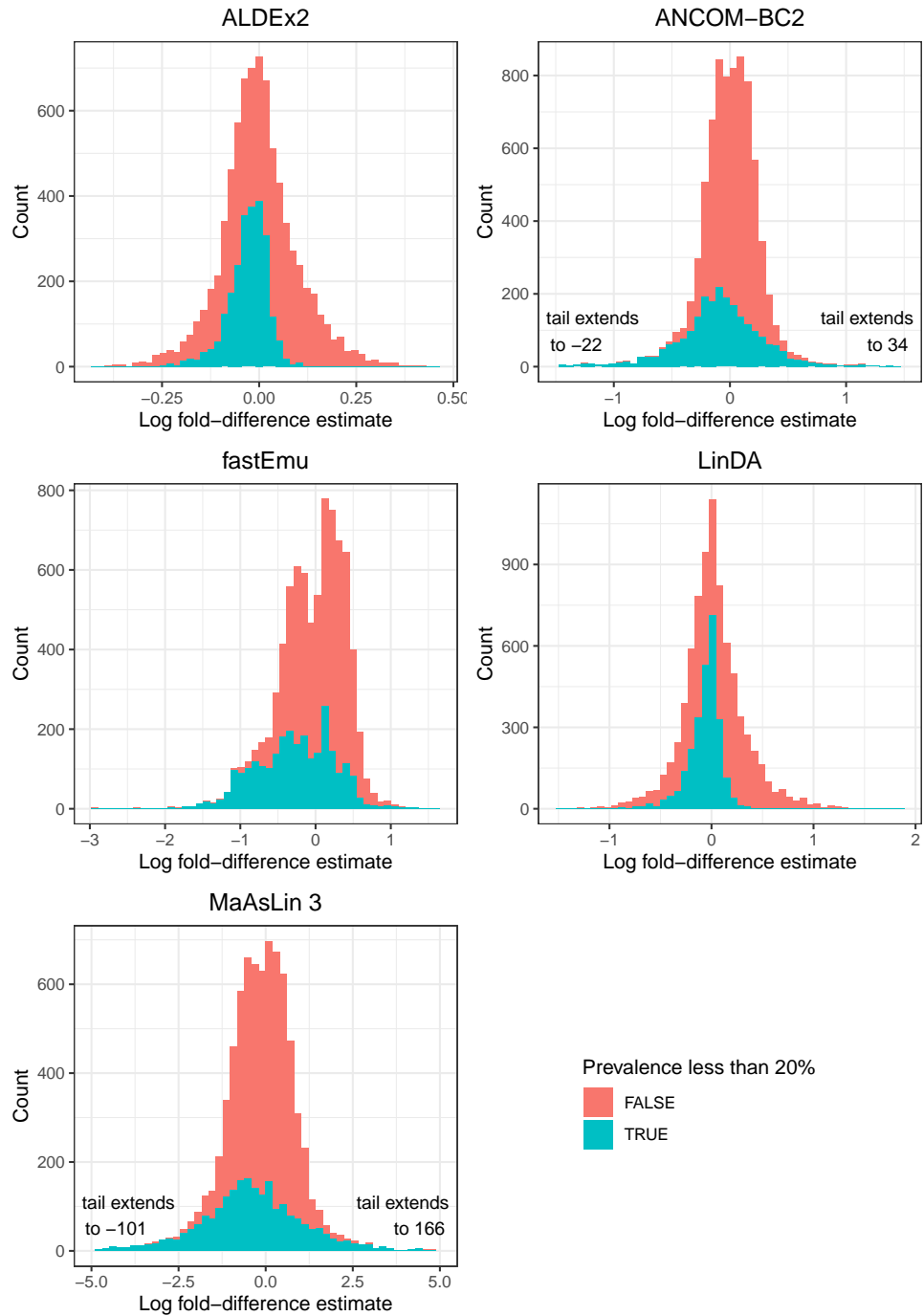

Fig 4: Histograms of log fold-difference in abundance estimates associated with a one-unit increase in temperature. Log fold-differences are estimated for 8,041 taxa. Bars are filled based on the proportion of taxa that are detected in fewer than 20% of samples. Each histogram represents a differential abundance method.

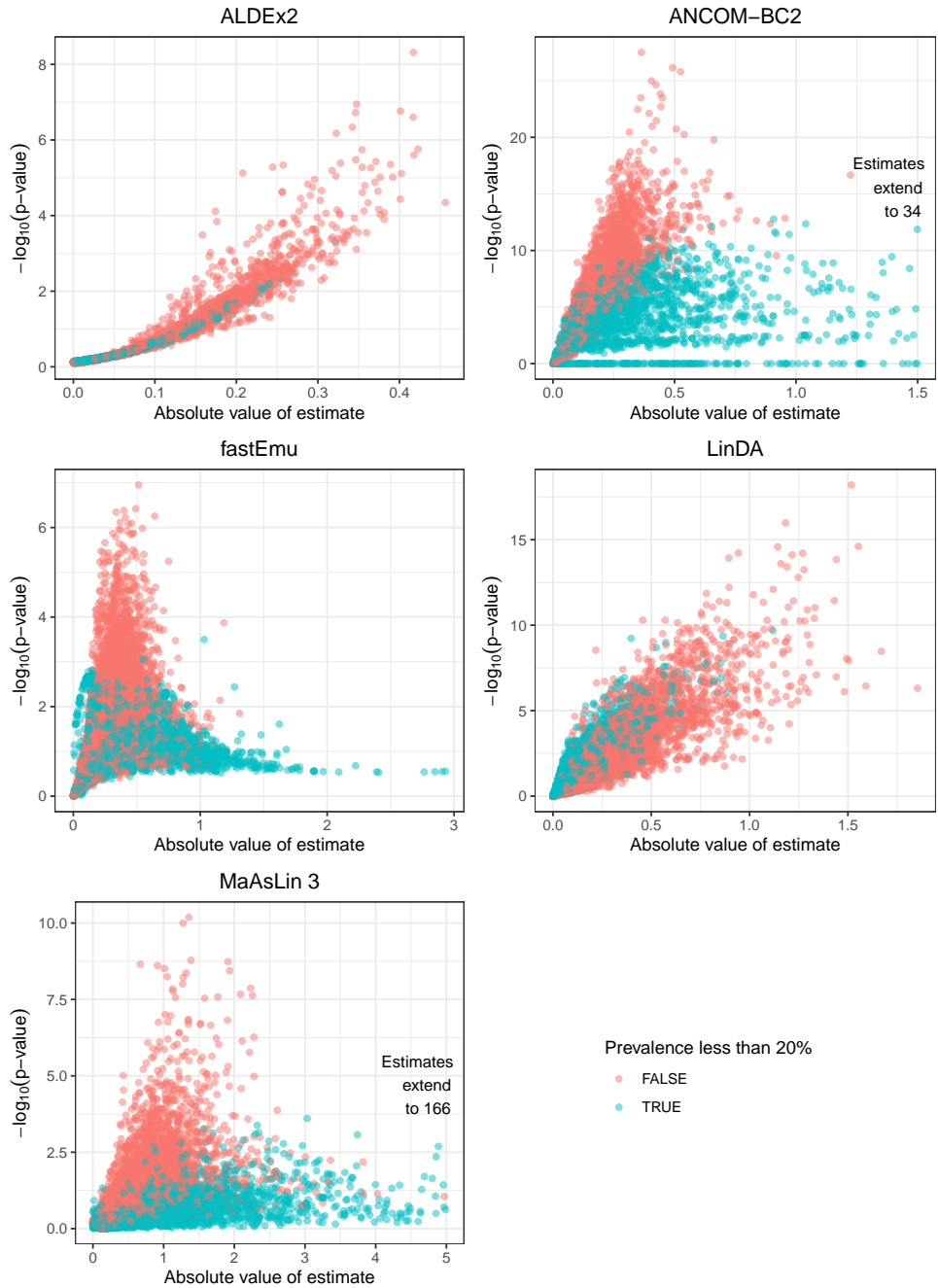

Fig 5: Scatterplots of absolute value of log fold-difference estimates by $-\log_{10}$ p-values. Each point represents one of the 8,041 taxa, and is colored based on whether or not the taxon is detected in fewer than 20% of samples. Each plot represents a differential abundance method.

**3.3. Comparison of inference.** For all taxa that appear only in one or two samples, ANCOM-BC2 assigns p-values of 1 and MaAsLin3 assigns missing p-values. The 285 taxa that have missing estimates from ALDEx2 also have missing p-values. We use p-values from ALDEx2, ANCOM-BC2, *fastEmu*, LinDA, and MaAsLin3 to generate a set of q-values from each method (Storey, 2002) (A q-value is the minimum false discovery rate at which a given test’s null hypothesis is rejected; unlike a p-value, which is defined for a single test in isolation, a q-value is assigned to each test but is estimated based on a collection of tests performed.). We find that out of 8,041 taxa, the number of taxa with q-values less than 5% are 110 (ALDEx2), 6195 (ANCOM-BC2), 7222 (*fastEmu*), 4738 (LinDA) and 1120 (MaAsLin3). It is unsurprising that such a large proportion of taxa have small p-values and q-values across several of these methods, because we expect that there are few taxa that have no association between abundance and ocean temperature. For the 5% of taxa for which we run *radEmu*, *radEmu* and *fastEmu* produce similar p-values, with a Spearman correlation of 0.99.

In Figure 5, we show the relationship between the parameter estimates and transformed p-values for each method. There is a strong association between the parameter estimates and p-values for ALDEx2 and LinDA. In contrast, the estimates and p-values are less strongly correlated for ANCOM-BC2, *fastEmu*, and MaAsLin3. They each include a long tail of estimates with larger magnitudes (especially for ANCOM-BC and MaAsLin3) and larger p-values, indicating that the best guess is that these taxa have large effect sizes, but there is uncertainty about whether they are truly non-zero. The taxa with the smallest p-values from ANCOM-BC2, *fastEmu*, and MaAsLin3 are taxa with more moderate estimate magnitudes and higher prevalences. It is not typical to obtain strongly correlated estimates and p-values, as these values provide distinct information: the former about the size of the effect, and the latter about the strength of evidence against a null effect.

**3.4. Differential abundance analysis.** We will now investigate the differential abundance analysis results from *fastEmu*. We will focus on taxa that have large log fold-difference estimates and small q-values, as these are biologically relevant fold-differences for which there is strong evidence against a null effect. We will consider the 3,590 taxa with q-values less than 0.01, which lets us control the false discovery rate at the 1% level.

The 20 taxa with the largest effect sizes estimates amongst the $q < 0.01$ subset are shown in Table 1. These taxa span diverse marine bacterial and archaeal taxa. These include *Pelagibacter* bacteria and *Marine Group II archaea*, both of which include clades that are known to inhabit distinct thermal and geographic niches (Giovannoni, 2017; Freel et al., 2025; Tully, 2019).

**4. Data-driven reference set selection.** In this section we explore the impact of using the data and initial parameter estimates $\hat{\beta}_{kj} - s_{[J]}(\hat{\beta}_{k\cdot})$ to determine the reference set $S_k$. We discuss potential concerns with this approach, propose alternate approaches that use distinct portions of the data for reference set selection and for hypothesis testing, and compare these approaches through simulation.

Step 2 in the *fastEmu* algorithm (Section 1.2) involves using the estimated *radEmu* parameters $\hat{\beta}_{jk} - s_{[J]}(\hat{\beta}_{k\cdot})$ to identify a “reference set”: taxa that have log fold-differences (in covariate $k$) that are the closest to the smoothed median of all log fold-differences. Then, this reference set is used to define the parameter that is estimated and tested in steps 3 and 4 of the algorithm. This data-driven reference set selection approach is a clear case of “double dipping”, because the same data are used for two separate tasks in an analysis. Double dipping causes invalid inference in many settings, and has inspired the field of selective inference, which involves the study of how to rigorously “measure the strength of the resulting selections” after “searching through the data for the strongest associations” (Taylor and Tibshirani,

| Estimate | Prevalence | Lowest temperature | Highest temperature | Phylum | Genus | Species |
| --- | --- | --- | --- | --- | --- | --- |
| -1.62 | 0.11 | 11.10 | 18.12 | Bacteroidota | MAG-121220-bin8 | None |
| -1.31 | 0.21 | 11.10 | 18.83 | Pseudomonadota | None | None |
| -1.27 | 0.11 | 11.10 | 18.12 | Pseudomonadota | Pseudothioglobus | Pseudothioglobus singularis |
| 1.19 | 0.26 | 23.37 | 30.59 | Pseudomonadota | AG-339-G14 | AG-339-G14 sp902536295 |
| -1.16 | 0.40 | 11.10 | 30.59 | Pseudomonadota | Pelagibacter | None |
| -1.09 | 0.21 | 11.10 | 27.17 | Pseudomonadota | MED-G78 | MED-G78 sp902509965 |
| 1.08 | 0.25 | 21.76 | 30.59 | Marinisomatota | Marinisoma | None |
| 1.06 | 0.32 | 19.31 | 30.59 | Pseudomonadota | MED-G82 | MED-G82 sp902600705 |
| 1.05 | 0.30 | 21.76 | 30.59 | Thermoplasmatota | MGIIb-N1 | None |
| 1.04 | 0.16 | 25.03 | 30.59 | Verrucomicrobiota | MB11C04 | MB11C04 sp002694035 |
| -1.03 | 0.18 | 11.10 | 19.25 | Thermoplasmatota | MGIIb-O2 | None |
| 1.02 | 0.21 | 19.77 | 30.59 | Pseudomonadota | TMED112 | None |
| 1.02 | 0.19 | 22.24 | 30.59 | None | None | None |
| 0.99 | 0.21 | 20.48 | 30.59 | Thermoplasmatota | MGIIa-L3 | None |
| -0.98 | 0.23 | 11.10 | 21.49 | Pseudomonadota | MED-G09 | MED-G09 sp002704345 |
| -0.96 | 0.21 | 11.10 | 27.17 | Actinomycetota | Actinomarina | None |
| 0.95 | 0.26 | 22.24 | 30.59 | Marinisomatota | GCA-002708715 | None |
| -0.95 | 0.21 | 11.10 | 21.49 | Thermoproteota | Nitrosopumilus | None |
| 0.94 | 0.23 | 18.12 | 30.59 | Pseudomonadota | None | None |
| -0.93 | 0.37 | 11.10 | 27.64 | Pseudomonadota | Pelagibacter | None |

TABLE 1

A set of 20 taxa with *fastEmu* $q$-values less than 0.01 and the largest effect size magnitudes. The effect size magnitudes range from 0.93 to 1.62, which correspond to fold-differences of 2.53 to 5.05.

2015). However, the major difference between the problems tackled in selective inference and our suggestion is that for most selective inference tasks, the first step is using the data to find the strongest associations and the second step is testing those associations. In our setting, we are instead searching for taxa with the *least* interesting association to inform a measure of center. We then use this baseline as a way to define a meaningful estimand, and then proceed to test hypotheses for all prespecified combinations of taxa and covariates.

A major concern about double dipping is losing Type I error rate control for a test that would otherwise be valid. An additional concern with using the reference set $S_k$ for parameter definition and then running robust score tests is that the parameter depends on the data, and therefore the test is not testing a well-defined null hypothesis. We address both of these concerns by showing that the distribution of robust score test statistics constructed using the full data approximates the distribution of robust score test statistics constructed using a reduced-taxon dataset. We prove these results formally for large sample sizes in Section 5, and demonstrate that this holds in finite samples via simulations in Section 4.2.

We expect that some researchers will still be wary of double-dipping in this context. We therefore also implement and study an alternative approach to choosing a reference set: splitting the data and using one split to choose the reference set and the other set for estimation and inference. We consider both sample-splitting and Poisson thinning (Neufeld et al., 2024a) as methods to split our data into two sets.

**4.1. Methods to split data for reference set selection and analysis.** In sample splitting, we randomly choose a subset of samples of size $n_r = \lceil \epsilon \times n \rceil$ for $\epsilon \in (0, 1)$ to use for reference set selection, and then use the remaining $n_a = n - n_r$ samples for estimation and inference. The $\epsilon$ value can be tuned to determine how much data to use for reference set selection versus analysis. This results in reference selection covariate data $X^r \in \mathbb{R}^{n_r \times p}$ and outcome data $Y^r \in \mathbb{R}^{n_r \times J}$, and analysis covariate data $X^a \in \mathbb{R}^{n_a \times p}$ and outcome data $Y^a \in \mathbb{R}^{n_a \times J}$. The benefit of sample splitting is that it doesn't require distributional assumptions on the outcome data.

In Poisson thinning, we separate each outcome $Y_{ij}$ into two counts $Y_{ij}^r$ and $Y_{ij}^a$ such that $Y_{ij}^r + Y_{ij}^a = Y_{ij}$. We use $Y^r \in \mathbb{R}^{n \times p}$ for reference set selection and $Y^a \in \mathbb{R}^{n \times J}$, and use $X$ as covariate data for both tasks. If the values $Y_{ij}$ are truly distributed as Poisson random variables, then the resulting $Y_{ij}^r$ and $Y_{ij}^a$ will also be distributed as Poisson, will be independent, and will have $\mathbb{E}[Y_{ij}^a] = \epsilon \times \mathbb{E}[Y_{ij}]$ and $\mathbb{E}[Y_{ij}^r] = (1 - \epsilon) \times \mathbb{E}[Y_{ij}]$ for $\epsilon \in (0, 1)$, where $\epsilon$ is a tuning parameter to determine what approximate proportion of each count should be included in $Y^r$. However, if the counts are not Poisson distributed, then the two datasets will not be fully independent and $Y_{ij}^r$ and $Y_{ij}^a$ will not be Poisson distributed. For example, Neufeld et al. (2024b) show that when applying Poisson thinning to data that is generated according to a negative binomial distribution with overdispersion parameter $b$, there will be a positive correlation between the $Y_{ij}^r$ and $Y_{ij}^a$ for all $i$ and $j$, and the correlation will be a function of $\epsilon$, $\mu_{ij}$, and $b$. This correlation increases as the overdispersion in the data increases. One could also consider thinning with other distributions, but each will require its own parametric assumptions.

When using sample splitting or Poisson thinning to produce two separate (and ideally independent) datasets, we will use the first set $Y^r$ to choose our reference sets. We will do this using the `fastEmu` algorithm, following steps 1 and 2 of the algorithm described in Section 1.2. We will refer to the reference sets from sample splitting as $S_k^{ss}$ and the reference sets from Poisson thinning as $S_k^{th}$. Once we have a reference set, we will estimate parameters and perform robust score tests using $Y^a$.

While the clear benefit of using sample splitting or thinning is that they provide separate reference set selection and analysis datasets, both approaches also have several drawbacks. Both sample splitting and thinning are random procedures and repeated analyses could lead to different reference sets and analysis results. Sample splitting is also associated with a loss in power, because we will have fewer observations to use in our analysis dataset. Poisson thinning cannot be directly applied to a non-integer dataset because only integers can be thinned with the Poisson distribution. Therefore, while this approach may be appropriate for amplicon data analysis, it may not be appropriate for many metagenomic analyses where coverages take on non-integer values. In practice, Poisson thinning can be applied to non-integer $Y_{ij}$ values by first rounding all $Y_{ij}$ values to the nearest integer.

**4.2. Reference set selection simulations.** In this section we perform additional Type I error and power simulations to compare the performance of `fastEmu` using the data-driven reference sets to `fastEmu` using the reference sets constructed using sample splitting and thinning. We also include `radEmu`, which implicitly uses a reference set $\{1, \dots, J\} := [J]$, and `fastEmu` using the true reference sets $S_k^*$ as comparison methods. In these simulations, $s_{[J]}(\beta_{k\cdot}) = s_{S_k^*}(\beta_{k\cdot})$ for all $k$ by construction.

**4.2.1. Simulation set-up.** We set $J = 250$ and choose a reference set size of $m = 25$. We simulate data with sample sizes of $n \in \{25, 50, 250\}$, and a balanced design matrix $X_{i0} = 1$ and $X_{i1} \in \{0, 1\}$. For each $J$, we construct a $\beta \in \mathbb{R}^{2 \times J}$ matrix. Again we set the first row of $\beta$ (corresponding to the intercept column of the design matrix) such that the odd elements are an evenly spaced sequence of length $J/2$ from $-3$ to $3$ and the even elements are an evenly spaced sequence of length $J/2$ from $3$ to $-3$. We set the second row of $\beta$ (corresponding to the covariate column of the design matrix) to be $5 \times \sinh(x)/\sinh(10)$, in which $x$ is defined as a linearly increasing sequence from $-10$ to $10$ with length $J$. In the Type I error rate simulations, we set $\beta_{k^*, j^*} = 0$ for $k^* = 1$ and $j^* = J/2$. In the power simulations, we set $\beta_{k^*, j^*} = b$ for $b \in \{0.5, 1, 2, 3, 4, 5\}$. This $J$ is relatively small and the setup is less complex compared to earlier simulations (Section 2) in order to more transparently contrast the options for reference set construction.

In the Type I error rate simulations, we generate data under the null hypothesis that $\beta_{k^*j^*} - s_{[J]}(\beta_{k^*}) = 0$ and also by construction $\beta_{k^*j^*} - s_{S_k^*}(\beta_{k^*}) = 0$ for $k^* = 1$ and $j^* = J/2$. In this framework, we will simultaneously evaluate the ability of each approach to reject this null hypothesis at a nominal level, regardless of whether the approach estimates and tests parameters of the form $\beta_{jk} - s_{[J]}(\beta_k)$ (the `radEmu` approach) or $\beta_{jk} - s_{S_k^*}(\beta_k)$ (each of the `fastEmu` approaches).

Each time that simulated data are generated, we draw $n$ values independently from a Normal distribution to use as the $z$ vector. Once we have $X$, $\beta$, and $z$, we can compute means of the form $\mu_{ij} = \exp(X_i\beta^j + z_i)$ for $i \in \{1, \dots, n\}$ and $j \in \{1, \dots, J\}$. We draw each $Y_{ij}$ value independently from either a Poisson distribution or a zero-inflated negative binomial (ZINB) distribution. The Poisson distribution is fully parameterized by the mean model. In order to draw from the ZINB distribution, we first draw from a Negative Binomial distribution with mean $\mu_{ij}/0.4$ and dispersion parameter $\phi = 5/(0.4^2)$ (this has a variance of $\mu_{ij}/0.4 + \mu_{ij}^2/5$). We then multiply this with an independent draw from a Bernoulli distribution with probability of success 0.4. This corresponds to a set of data with roughly 60% zeroes.

For both sample splitting and thinning we set the tuning parameter $\epsilon = 0.25$. When using the sample splitting approach, we assign $\lceil 0.25 \times n \rceil = 13$ samples to the reference set selection dataset ($X^r \in \mathbb{R}^{13 \times 2}$, $Y^r \in \mathbb{R}^{13 \times 250}$) and the remaining $\lfloor 0.75 \times n \rfloor = 37$ samples to the analysis dataset ($X^a \in \mathbb{R}^{37 \times 2}$, $Y^a \in \mathbb{R}^{37 \times 250}$). When using the Poisson thinning approach, we generate $Y^r \in \mathbb{R}^{50 \times 250}$ and $Y^a \in \mathbb{R}^{50 \times 250}$ such that $Y_{ij}^r + Y_{ij}^a = Y_{ij}$ and $\mathbb{E}[Y_{ij}^a] = 0.25 \times \mathbb{E}[Y_{ij}]$ and $\mathbb{E}[Y_{ij}^r] = 0.75 \times \mathbb{E}[Y_{ij}]$. We generate outcome data using a Poisson model (in which Poisson thinning leads to independent reference set selection and analysis sets) and using the ZINB approach used in Section 2. For simulation settings with Poisson distributed data, $Y_{ij}^r$ and $Y_{ij}^a$ are also Poisson distributed. For simulations settings with ZINB data, the expectations $0.25 \times \mathbb{E}[Y_{ij}]$ and $\mathbb{E}[Y_{ij}^a] = 0.75 \times \mathbb{E}[Y_{ij}]$ still hold, but we will not have results about the distribution of $Y_{ij}^r$ and $Y_{ij}^a$, and the two datasets will be positively correlated (Neufeld et al., 2024b).

**4.2.2. Simulation results.** Quantile-quantile plots of p-values from this simulation can be seen in Figure 6. The empirical p-value distributions are nearly the same across all five reference set approaches. For each of these simulation settings, the distribution of p-values under the equivalent null hypotheses $\beta_{k^*j^*} - s_{[J]}(\beta_{k^*}) = 0$ and $\beta_{k^*j^*} - s_{S_k^*}(\beta_{k^*}) = 0$ for $k^* = 1$ and $j^* = J/2$ are approximately Uniform(0, 1). Therefore, in this simulation we do not see a loss of Type I error rate control when using data-driven reference set $S_k$ directly from the data or sets $S_k^{ss}$ and $S_k^{th}$, when compared to pre-specified reference sets $[J]$ and $S_k^*$.

The results from the power simulations can be seen in Figure 7. In the Poisson settings, the power is nearly the same for all approaches, although it is slightly lower for the sample splitting and thinning approaches for signals with $\beta_1 < 1$. The same pattern holds for ZINB data with $n = 250$. However, for the ZINB data with $n \in \{25, 50\}$, the power for tests with reference sets $S_k$ and $S_k^*$ is higher than the power for the test with reference set $[J]$, especially for medium to large signals. We hypothesize that the difference in power can be explained by a higher estimated standard error of the score vector when we construct a reference set with all taxa $[J]$, versus a smaller subset of taxa in $S_k$ and $S_k^*$.

The mean squared errors (MSEs) for estimating the parameter $\beta_{k^*j^*} - s_{[J]}(\beta_{k^*})$ in these Type I error simulations are contained in Table 2. Across all simulation settings, the estimation MSE is very similar between the approaches that use reference sets $[J]$, $S_k$ and $S_k^*$, all of which have a lower error than the approaches that use reference sets $S_k^{ss}$ and $S_k^{th}$.

These simulations demonstrate that Type I error is not inflated by using a data-driven method to select a reference set, including by using all the data to select the set, using sample splitting or sample thinning. However, the sample splitting and thinning approaches are associated with lower power, which is more noticeable for the ZINB data with smaller sample sizes. The lower power makes sense for sample splitting, as analysis is done using a smaller sample size than the other constraints. However, the cause of reduced power for the thinning approaches is not obvious, because testing error should not be affected by rescaling. We also find that for sparse ZINB data and smaller sample sizes, the approach that uses reference sets

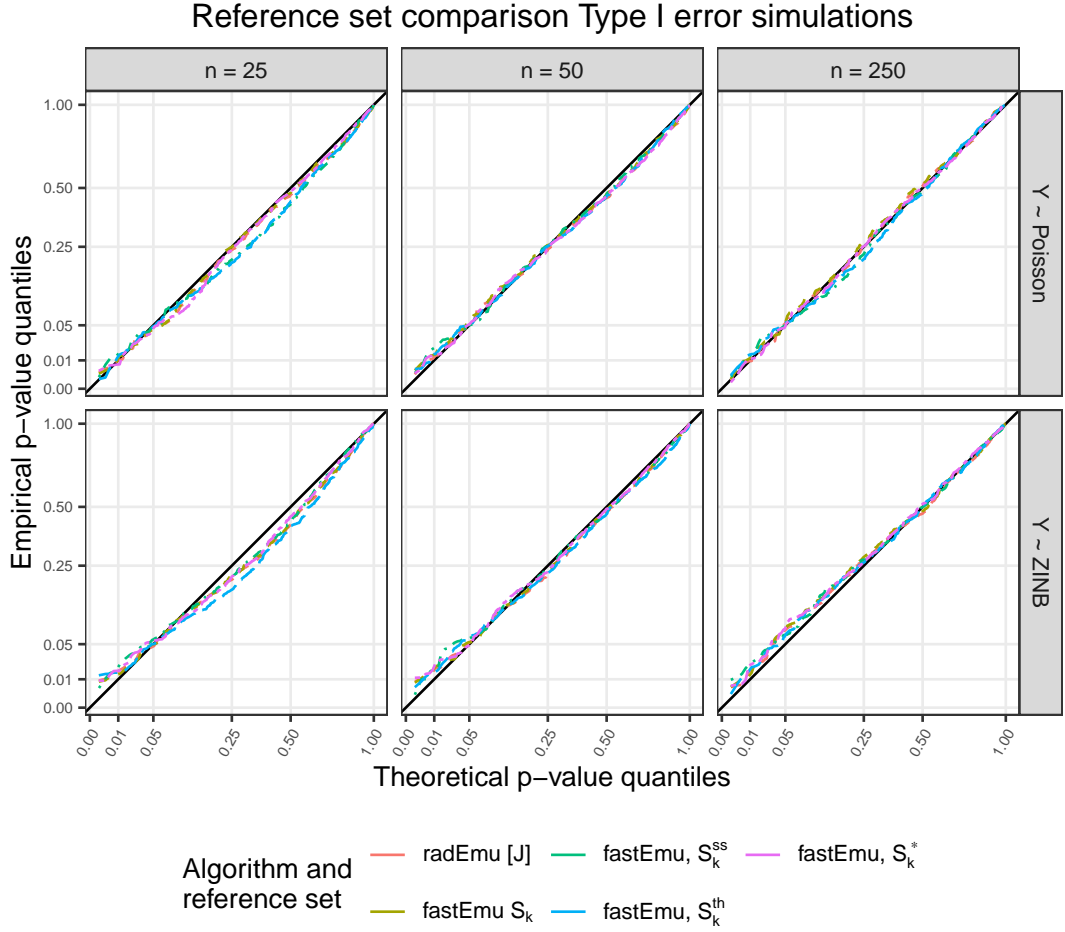

Fig 6: Quantiles of p-values obtained from the reference set Type I error rate simulation compared to quantiles of a $\text{Uniform}(0, 1)$ distribution. Data is simulated under the null hypothesis $\beta_{k^*j^*} - s_{[J]}(\beta_{k^*}) = 0$. P-values are generated from robust score tests using reference sets $[J]$ (the `radEmu` approach), $S_k$ (the `fastEmu` approach described in Section 1.2), $S_k^*$ (the true unknown reference set), $S_k^{ss}$ (a reference set determined using sample splitting), and $S_k^{th}$ (a reference set determined using Poisson thinning). The $x = y$ line is shown in black, and represents quantiles of p-values from a test that controls Type I error rate at a nominal rate across the full range of p-values. A conservative test will produce a curve above the line $x = y$ for small p-values and an anti-conservative test will produce a curve below it. Results based on 500 trials.

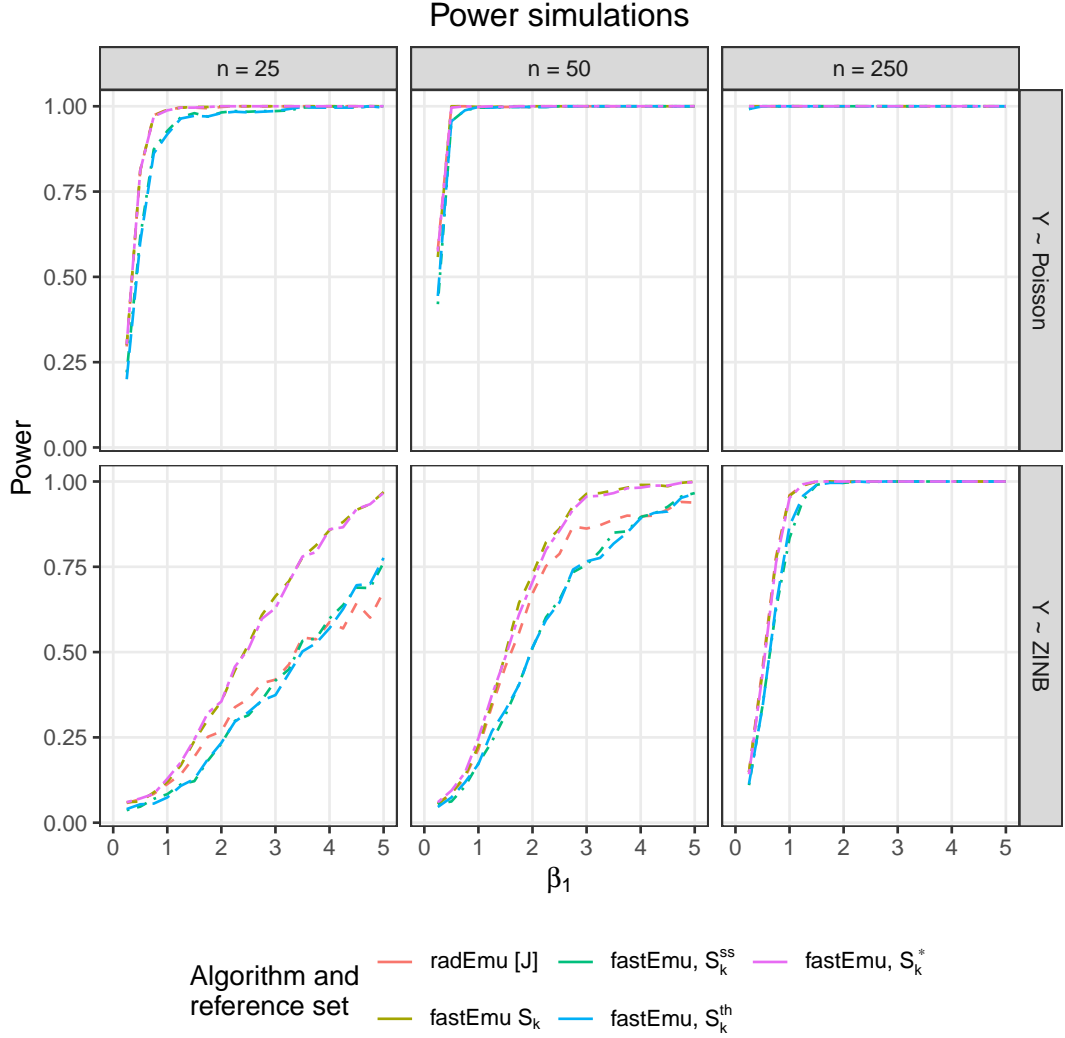

Fig 7: Power curve plots using p-values from reference set power simulations. Data is simulated under the 20 different specific alternatives based on different magnitudes of $\beta_{k^*j^*} - s_{[J]}(\beta_{k^*})$. P-values are generated from robust score tests using reference sets $[J]$ (the `radEmu` approach), $S_k$ (the `fastEmu` approach described in Section 1.2), $S_k^*$ (the true unknown reference set), $S_k^{ss}$ (a reference set determined using sample splitting), and $S_k^{th}$ (a reference set determined using Poisson thinning).

$S_k$ and $S_k^*$ lead to a test with higher power than the `radEmu` approach using the full set of taxa $[J]$. Therefore, from a pragmatic viewpoint, we recommend using the data-driven reference set from the whole dataset. However, if this double dipping is of concern to the user of our method, we suggest using sample splitting or thinning to choose a reference set.

### 5. Technical results and proofs. In this section, we will prove three results.

1. <u>`fastEmu` and `radEmu` target the same parameters when reference sets are prespecified.</u> For a prespecified reference set $S_R$, the identifiable estimands defined by the `fastEmu` model and algorithm coincide with those defined by the `radEmu` model algorithm on the retained categories $j \in S_R$ (Propositions 1 and 2).

| Sample size | Data | Set $[J]$ | Set $S_k$ | Set $S_k^*$ | Set $S_k^{ss}$ | Set $S_k^{th}$ |
| --- | --- | --- | --- | --- | --- | --- |
| 25 | Poisson | 0.025 | 0.025 | 0.024 | 0.040 | 0.041 |
| 50 | Poisson | 0.011 | 0.011 | 0.011 | 0.015 | 0.016 |
| 250 | Poisson | 0.002 | 0.002 | 0.002 | 0.003 | 0.003 |
| 25 | ZINB | 1.061 | 1.061 | 1.008 | 1.954 | 1.843 |
| 50 | ZINB | 0.363 | 0.365 | 0.348 | 0.555 | 0.526 |
| 250 | ZINB | 0.069 | 0.069 | 0.067 | 0.091 | 0.090 |

TABLE 2

MSE from estimating $\beta_{k*}j^* - s_{[J]}(\beta_{k*})$ in Type 1 error simulations from approaches with different reference sets. These results are aggregated across 500 trials.

2. The `fastEmu` test statistic has the correct null distribution when the reference set is pre-specified. Under the target null hypothesis $\beta_{kj'} - s_{S_R}(\beta_{k*}) = 0$ with prespecified reference set $S_R$, the robust score statistic constructed with the algorithm described in Section 1.2 converges to a $\chi_1^2$ distribution (Proposition 3).
3. Using a data-driven reference set has negligible asymptotic impact. The data-driven reference set $S_k$ equals the (unknown) oracle reference set $S_k^*$ with probability tending to one (Proposition 4), and consequently the data-driven test statistic shares the $\chi_1^2$ asymptotic distribution (Proposition 5).

5.1. *Setup and notation.* We observe independent pairs $(X_i, Y_i)$, $i \in [n] := \{1, \dots, n\}$, where $X_i \in \mathbb{R}^{1 \times p}$ is a row of the design matrix $\mathbf{X} \in \mathbb{R}^{n \times p}$ (with intercept, i.e. $X_{i0} = 1$) and $Y_i = (Y_{i1}, \dots, Y_{iJ}) \in \mathbb{R}_{\geq 0}^J$ is a vector of nonnegative outcomes over $J$ categories (taxa). We write $[J] := \{1, \dots, J\}$. Covariates are indexed by $k \in \{0, 1, \dots, p-1\}$ and taxa by $j \in [J]$. For a matrix $\beta \in \mathbb{R}^{p \times J}$ we write $\beta_{\cdot j} \in \mathbb{R}^{p \times 1}$ for its $j$-th column, $\beta_{k\cdot} \in \mathbb{R}^{1 \times J}$ for its $k$-th row, and $\beta_{kj}$ for its $(k, j)$ entry. Throughout, $\mathbf{X}$ has full column rank (an assumption maintained throughout Clausen, Teichman and Willis 2024).

5.2. *The full and reduced models.* Following Clausen, Teichman and Willis (2024), the `radEmu` (“full”) mean model posits sample-specific nuisance effects $z_1, \dots, z_n \in \mathbb{R}$ and $\beta \in \mathbb{R}^{p \times J}$ such that

$$(M_F) \quad \log \mathbb{E}[Y_{ij} \mid X_i, z_i] = z_i + X_i \beta_{\cdot j}, \quad i \in [n], j \in [J].$$

As shown by Clausen, Teichman and Willis (2024), model $(M_F)$ is identifiable only up to the transformation $\beta_{\cdot j} \mapsto \beta_{\cdot j} + c$, $z_i \mapsto z_i - X_i c$ (for $c \in \mathbb{R}^p$ common to all $j$); consequently $\beta$ itself is not identifiable, but any *location-equivariant contrast* of a row of $\beta$ is. Our estimands are of this form: for a centering (identifiability) function $g: \mathbb{R}^J \rightarrow \mathbb{R}$ satisfying

$$(3) \quad g(v + a \mathbf{1}_J) = g(v) + a \quad \text{for all } v \in \mathbb{R}^J, a \in \mathbb{R},$$

the quantity $\beta_{kj} - g(\beta_{k\cdot})$ is invariant to the transformation above and is therefore identifiable.

`fastEmu` works with a subset of taxa. Fix a nonempty set $S_R \subseteq [J]$ (this could be $S_k$, as described in Section 1.2), and let $Y^\dagger = (Y_{ij})_{i \in [n], j \in S_R}$ be the $n \times |S_R|$ submatrix of $\mathbf{Y}$ retaining the columns in $S_R$. `fastEmu` fits a `radEmu` (“reduced”) model to $Y^\dagger$: there exist $z_1^\dagger, \dots, z_n^\dagger \in \mathbb{R}$ and $\beta^\dagger \in \mathbb{R}^{p \times |S_R|}$ (columns indexed by $S_R$) such that

$$(M_R) \quad \log \mathbb{E}[Y_{ij} \mid X_i, z_i^\dagger] = z_i^\dagger + X_i \beta_{\cdot j}^\dagger, \quad i \in [n], j \in S_R.$$

Throughout we assume that both the full model $(M_F)$ and the reduced model $(M_R)$ are correctly specified.

We brief note two facts about the smoothed median: (i) $s_S(b)$ depends on $b$ only through $\{b_j : j \in S\}$; and (ii) $s_S(b + a \mathbf{1}) = s_S(b) + a$ for all $a \in \mathbb{R}$.

5.3. *The robust score statistic.* To test a hypothesis of the form $\beta_{k'j'} - s_{Sr}(\beta_k)$, for reference sets of the form $Sr = [J]$ (as in `radEmu`) or $Sr = S_k$ (as in `fastEmu`), `radEmu` uses the misspecification-robust score statistic of White (1982),

$$(4) \quad T_{RS} = S_{H_0}^T I_{H_0}^{-1} F_{H_0}^T (F_{H_0} I_{H_0}^{-1} D_{H_0} I_{H_0}^{-1} F_{H_0}^T)^{-1} F_{H_0} I_{H_0}^{-1} S_{H_0},$$

where $S_{H_0}$ is the score evaluated at the maximum likelihood estimate under the null, $I_{H_0}$ is a consistent estimate of the information matrix under the null, $D_{H_0} = \sum_{i=1}^n s_i s_i^T$ is the empirical outer product of the per-observation score contributions $s_i$, and $F_{H_0} = \partial h / \partial \beta^T$ evaluated at the null estimate where $h(\beta) = \beta_{k'j'} - s_{Sr}(\beta_k)$. In practice `radEmu` applies the small-sample correction of Guo et al. (2005), replacing $T_{RS}$ by $\frac{n}{n-1} T_{RS}$; since $n/(n-1) \rightarrow 1$, this factor does not affect any asymptotic distribution and we suppress it below. When (4) is computed from the full model ($M_F$) we denote it $T$, and when computed from the reduced model ($M_R$) we denote it $T^\dagger$.

5.4. *Parameter equivalence between `radEmu` and `fastEmu` with predetermined reference sets.* In this section, we will prove that for a predetermined reference set, the identifiable estimands targeted by `radEmu` and `fastEmu` are equivalent. Define $Sr$ as a predetermined reference set for covariate $k$. We first show this for contrasts against a common reference taxon, and then extend to centering functions generally.

PROPOSITION 1. Assume ($M_F$) and ($M_R$) hold and $\mathbf{X}$ has full column rank. Then for every pair $j, j' \in S_R$ and every $k \in \{0, \dots, p-1\}$,

$$(5) \quad \beta_{kj} - \beta_{kj'} = \beta_{kj}^\dagger - \beta_{kj'}^\dagger.$$

PROOF. Fix $j, j' \in S_R$. Differencing the mean model ($M_F$) across the two taxa eliminates the sample effect $z_i$: for every $i \in [n]$,

$$(6) \quad \log \mathbb{E}[Y_{ij} | X_i] - \log \mathbb{E}[Y_{ij'} | X_i] = X_i \beta_{.j} - X_i \beta_{.j'} = X_i (\beta_{.j} - \beta_{.j'}).$$

Because $j, j' \in S_R$, the reduced model ($M_R$) applies to both taxa, and differencing it likewise eliminates $z_i^\dagger$:

$$(7) \quad \log \mathbb{E}[Y_{ij} | X_i] - \log \mathbb{E}[Y_{ij'} | X_i] = X_i (\beta_{.j}^\dagger - \beta_{.j'}^\dagger).$$

The left-hand sides of (6) and (7) are identical — they are log-ratios of the *same* conditional means of the observed outcomes — so their right-hand sides agree: $X_i (\beta_{.j} - \beta_{.j'}) = X_i (\beta_{.j}^\dagger - \beta_{.j'}^\dagger)$ for every $i \in [n]$. Stacking these $n$ scalar identities gives

$$\mathbf{X} (\beta_{.j} - \beta_{.j'} - \beta_{.j}^\dagger + \beta_{.j'}^\dagger) = \mathbf{0}_n.$$

Since $\mathbf{X}$ has full column rank, the only vector in its kernel is $\mathbf{0}_p$, hence $\beta_{.j} - \beta_{.j'} = \beta_{.j}^\dagger - \beta_{.j'}^\dagger$. Reading off the $k$-th coordinate yields (5). □

We now generalize from a single reference taxon to the class of centering functions used by `fastEmu`. Let $v[S_R] \in \mathbb{R}^{|S_R|}$ denote the subvector of $v \in \mathbb{R}^J$ indexed by $S_R$. Given a full-model centering function $g_F : \mathbb{R}^J \rightarrow \mathbb{R}$ and a reduced-model centering function $g_R : \mathbb{R}^{|S_R|} \rightarrow \mathbb{R}$, define the compatible class

$$(8) \quad \mathcal{G}_{S_R} = \left\{ (g_F, g_R) : g_F(v) = g_R(v[S_R]) \text{ for all } v \in \mathbb{R}^J \right\}.$$

PROPOSITION 2. Let $g_F$ and $g_R$ each satisfy the location-equivariance property (3), and suppose $(g_F, g_R) \in \mathcal{G}_{S_R}$. Then under the assumptions of Proposition 1,

$$\beta_{kj} - g_F(\beta_k) = \beta_{kj}^\dagger - g_R(\beta_k^\dagger) \quad \text{for all } k \in \{0, \dots, p-1\} \text{ and all } j \in S_R.$$

PROOF. Fix $k$ and a reference index $r \in S_R$; write $c := \beta_{kr}$ and $c^\dagger := \beta_{kr}^\dagger$. By Proposition 1, for every $j \in S_R$,

$$(9) \quad \beta_{kj} - c = \beta_{kj}^\dagger - c^\dagger, \quad \text{equivalently} \quad (\beta_k - c \mathbf{1}_J)[S_R] = \beta_k^\dagger - c^\dagger \mathbf{1}_{|S_R|}.$$

Using location equivariance (3) of $g_F$ (with $a = -c$), the compatibility $g_F(v) = g_R(v[S_R])$ from (8) applied to $v = \beta_k - c \mathbf{1}_J$, the identity (9), and location equivariance of $g_R$ (with $a = -c^\dagger$),

$$\begin{aligned} \beta_{kj} - g_F(\beta_k) &= (\beta_{kj} - c) - (g_F(\beta_k) - c) \\ &= (\beta_{kj} - c) - g_F(\beta_k - c \mathbf{1}_J) && \text{[equivariance of } g_F\text{]} \\ &= (\beta_{kj} - c) - g_R((\beta_k - c \mathbf{1}_J)[S_R]) && [(g_F, g_R) \in \mathcal{G}_{S_R}] \\ &= (\beta_{kj}^\dagger - c^\dagger) - g_R(\beta_k^\dagger - c^\dagger \mathbf{1}_{|S_R|}) && \text{[by (9)]} \\ &= (\beta_{kj}^\dagger - c^\dagger) - (g_R(\beta_k^\dagger) - c^\dagger) && \text{[equivariance of } g_R\text{]} \\ &= \beta_{kj}^\dagger - g_R(\beta_k^\dagger). && \square \end{aligned}$$

5.5. Asymptotic distribution of test statistics generated using *fastEmu* testing procedure with prespecified reference sets. Proposition 2 shows that, for $k'$ and $j' \in S_R$, the full-model null

$$H_0: \beta_{k'j'} - s_{Sr}(\beta_{k'}) = 0$$

and the reduced-model null

$$H_0^\dagger: \beta_{k'j'}^\dagger - s_{Sr}(\beta_{k'}^\dagger) = 0$$

are equivalent: because $\beta_{kj} - s_{Sr}(\beta_k) = \beta_{kj}^\dagger - s_{Sr}(\beta_k^\dagger)$ for all $k$ and $j \in S_R$, one equals zero if and only if the other does. This equivalence justifies testing the computationally inexpensive $H_0^\dagger$ in place of $H_0$. We now show that the resulting test has the correct asymptotic null distribution.

Let $\hat{\beta}_{H_0^\dagger}^\dagger$ be the *radEmu* estimator of the reduced-model parameters under $(M_R)$ subject to the restriction $H_0^\dagger$, and let $T_{RS}^\dagger$ be the robust score statistic (4) evaluated at $\hat{\beta}_{H_0^\dagger}^\dagger$ with $h(\beta^\dagger) = \beta_{kj}^\dagger - s_{Sr}(\beta_k^\dagger)$.

PROPOSITION 3. Fix $k' \in \{1, \dots, p-1\}$ and $j' \in S_R$ for fixed $S_R$. Then under $H_0$ (equivalently, $H_0^\dagger$), $T_{RS}^\dagger \xrightarrow{d} \chi_1^2$.

PROOF. By Proposition 2, $H_0$ and $H_0^\dagger$ are equivalent, so it suffices to work under $H_0^\dagger$. The reduced model $(M_R)$ is a *radEmu* model fit to the $|S_R|$-category outcome $Y^\dagger$; the asymptotic theory of Clausen, Teichman and Willis (2024) therefore applies to $\hat{\beta}_{H_0^\dagger}^\dagger$. In particular, the reduced-model estimator is consistent and asymptotically normal, and $I_{H_0^\dagger}^\dagger$ and $D_{H_0^\dagger}^\dagger$ in (4) are consistent for the information matrix and the score covariance when defined using the reduced model $(M_R)$. Consequently $T_{RS}^\dagger$ is exactly the robust score statistic of White (1982) for a single restriction, evaluated at the constrained estimator. By Theorem 3.5 of White (1982), under $H_0^\dagger$, $T_{RS}^\dagger \xrightarrow{d} \chi_q^2$ with $q = 1$ degree of freedom due to $h(\beta) \in \mathbb{R}^1$. □

5.6. *Validity with data-driven reference sets.* Propositions 2 and 3 require a *pre-specified* reference set $S_R$. `fastEmu` instead selects the reference set from the data. In this section we show that this selection has no asymptotic cost: the data-driven reference set recovers the oracle reference set $S_k^*$ with probability tending to one, and the resulting test statistic retains the $\chi_1^2$ limit.

Fix a covariate index $k \in \{1, \dots, p-1\}$ throughout this section, and use the smoothed median constraint function. Define the true and estimated identifiable magnitudes

$$m_j := |\beta_{kj} - s_{[J]}(\beta_{k\cdot})|, \quad \hat{m}_j := |\hat{\beta}_{kj} - s_{[J]}(\hat{\beta}_{k\cdot})|, \quad j \in [J],$$

where $\hat{\beta}$ is the `radEmu` estimator fit under $H_A$. For a target reference-set size $J_r \in \{1, \dots, J\}$, the oracle (“known”) reference set and the data-driven reference set are

$$S_k^* := \{j \in [J] : \text{rank}(m_j) \leq J_r\}, \quad S_k := \{j \in [J] : \text{rank}(\hat{m}_j) \leq J_r\},$$

that is, the taxa with the $J_r$ smallest magnitudes. We assume that the true magnitudes $m_1, \dots, m_J$ are pairwise distinct and therefore $S_k^*$ is uniquely defined, and the gap $\Delta := \min_{j \neq j'} |m_j - m_{j'}|$ is strictly positive. We also use the consistency of the `radEmu` estimator of the identifiable parameter:

$$(10) \quad \hat{\beta}_{kj} - s_{[J]}(\hat{\beta}_{k\cdot}) \xrightarrow{p} \beta_{kj} - s_{[J]}(\beta_{k\cdot}) \quad \text{for each } j \in [J].$$

PROPOSITION 4. *Let ties among $\hat{m}_1, \dots, \hat{m}_J$, if any, be broken by an arbitrary rule. Then $\Pr(S_k = S_k^*) \rightarrow 1$ as $n \rightarrow \infty$.*

PROOF. The continuous mapping theorem gives $\hat{m}_j \xrightarrow{p} m_j$ for each $j \in [J]$. Since $J$ is fixed and finite, a union bound yields $M_n := \max_{j \in [J]} |\hat{m}_j - m_j| \xrightarrow{p} 0$. Let $j, j' \in [J]$ satisfy $m_j < m_{j'}$, so $m_{j'} - m_j \geq \Delta > 0$. On the event $\{M_n < \Delta/2\}$,

$$\hat{m}_{j'} \geq m_{j'} - M_n \geq m_j + \Delta - M_n > m_j + \Delta/2 > m_j + M_n \geq \hat{m}_j.$$

Hence, on $\{M_n < \Delta/2\}$, $\hat{m}_j < \hat{m}_{j'}$ whenever $m_j < m_{j'}$. Because the true magnitudes are pairwise distinct, the estimated magnitudes are then also pairwise distinct, so no tie-breaking is invoked. Their ordering also coincides with that of the true magnitudes, so $\text{rank}(\hat{m}_j) = \text{rank}(m_j)$ for every $j \in [J]$ and $S_k = S_k^*$. Therefore

$$\Pr(S_k = S_k^*) \geq \Pr(M_n < \Delta/2) \rightarrow 1.$$

□

Finally, we show that replacing the oracle reference set $S_k^*$ by the data-driven reference set $S_k$ does not change the *asymptotic* distribution of the test statistic. For a reference set $S \in \{S_k^*, S_k\}$ for covariate $k$ and the fixed target taxon $j'$, let $T_{RS}(S)$ and $T_{RS}^\dagger(S)$ denote the robust score statistics for testing $\beta_{kj'} - s_S(\beta_{k\cdot}) = 0$, computed respectively from the full model ($M_F$) on all $J$ taxa (using the smoothed median function on reference set $S$) and from the reduced model ($M_R$) fit to $Y^\dagger$ with retained columns $S_j := S \cup \{j\}$.

PROPOSITION 5. *Fix $k \in \{1, \dots, p-1\}$ and $j' \in [J]$. Under the null hypothesis*

$$H_0: \beta_{kj'} - s_{S_k^*}(\beta_{k\cdot}) = 0,$$

*and under the assumptions of Propositions 3 and 4,*

$$T_{RS}(S_k^*) \xrightarrow{d} \chi_1^2, \quad T_{RS}^\dagger(S_k^*) \xrightarrow{d} \chi_1^2, \quad T_{RS}(S_k) \xrightarrow{d} \chi_1^2, \quad T_{RS}^\dagger(S_k) \xrightarrow{d} \chi_1^2.$$

PROOF. The oracle reference set $S_k^*$ is a fixed subset of $[J]$, so the centering function $s_{S_k^*}$ is a pre-specified identifiability constraint. The convergence $T_{RS}(S_k^*) \xrightarrow{d} \chi_1^2$ under $H_0$ is therefore the standard `radEmu` robust score result for a pre-specified constraint. Then convergence $T_{RS}^\dagger(S_k^*) \xrightarrow{d} \chi_1^2$ follows from Proposition 3.

Each statistic is a fixed measurable function of the data and of the chosen reference set: write $T_{RS}(S) = \mathcal{T}(\text{data}, S)$ for a common map $\mathcal{T}$. The data-driven statistic evaluates this same map at the random set $S_k$, whereas the oracle statistic evaluates it at $S_k^*$. On the event $\{S_k = S_k^*\}$ the two arguments coincide, so $\mathcal{T}(\text{data}, S_k) = \mathcal{T}(\text{data}, S_k^*)$, i.e. $T_{RS}(S_k) = T_{RS}(S_k^*)$ pointwise. Hence, for any $\varepsilon > 0$,

$$\begin{aligned} \Pr(|T_{RS}(S_k) - T_{RS}(S_k^*)| \geq \varepsilon) &= \underbrace{\Pr(|T_{RS}(S_k) - T_{RS}(S_k^*)| \geq \varepsilon, S_k = S_k^*)}_{=0} + \\ &\quad \Pr(|T_{RS}(S_k) - T_{RS}(S_k^*)| \geq \varepsilon, S_k \neq S_k^*) \\ &\leq \Pr(S_k \neq S_k^*) \rightarrow 0 \end{aligned}$$

by Proposition 4. Therefore $T_{RS}(S_k) - T_{RS}(S_k^*) \xrightarrow{p} 0$. Then by Slutsky's theorem, $T_{RS}(S_k) \xrightarrow{d} \chi_1^2$. The same argument applies to the reduced statistics: on the event $\{S_k = S_k^*\}$ the retained column sets $S_j = S_k \cup \{j\}$ and $S_k^* \cup \{j\}$ coincide, so $Y^\dagger$ and hence $T_{RS}^\dagger(S_k) = T_{RS}^\dagger(S_k^*)$ coincide; thus $T_{RS}^\dagger(S_k) - T_{RS}^\dagger(S_k^*) \xrightarrow{p} 0$, and Slutsky's theorem together with $T_{RS}^\dagger(S_k^*) \xrightarrow{d} \chi_1^2$ gives $T_{RS}^\dagger(S_k) \xrightarrow{d} \chi_1^2$. □

Therefore, selecting the reference set from the same data used for testing therefore does not compromise asymptotic Type I error control. Moreover, because $S_k^*$ collects the taxa whose true log fold-differences are closest to the smoothed median, $s_{S_k^*}(\beta_{k\cdot}) \approx s_{[J]}(\beta_{k\cdot})$, and the tested null closely approximates the `radEmu` null $\beta_{kj'} - s_{[J]}(\beta_{k\cdot}) = 0$. This implies that when $|s_{S_k^*} - s_{[J]}|$ is small, `fastEmu` tests a hypothesis that is numerically very close to the `radEmu` null while substantially reducing computation.
